# State-dependent gating of sensory and cortical inputs by median raphe neurons

**DOI:** 10.64898/2026.11.13.766493

**Authors:** Wenqiang Huang, Dong V Wang

## Abstract

The mesopontine median raphe (MnR) plays an important role in regulating brain states, hippocampal network activity, and memory processes. Recent studies have revealed that the MnR comprises three major neuronal populations: serotonergic (5-HT), glutamatergic (Vglut3-expressing), and GABAergic neurons. How these distinct neuronal populations may gate sensory and cortical inputs across different brain states remains largely unknown. Here, we combined in vivo electrophysiological recording, sensory or optogenetic stimulation, and transgenic approaches to characterize the response properties of 5-HT, Vglut3, and GABAergic MnR neurons during wakefulness and sleep. We found that MnR neurons exhibited pronounced state-dependent responses to both sensory and cortical stimulation, with response amplitudes significantly greater during slow-wave sleep than during wakefulness. Auditory and somatosensory stimuli elicited fast responses in the MnR: 5-HT neurons were primarily suppressed, whereas Vglut3 and GABAergic neurons were primarily activated, suggesting the presence of direct inputs from sensory thalamic or brainstem structures. Retrograde tracing identified several brainstem structures and, notably, the medial prefrontal cortex as sources of direct projections to the MnR. Optogenetic stimulation of prefrontal inputs to the MnR evoked inhibition of 5-HT neurons but robust excitation of Vglut3 neurons during slow-wave sleep, suggesting top-down control of MnR activity by the prefrontal cortex. Together, these findings reveal that the MnR exhibits enhanced sensitivity to both ascending sensory and descending cortical inputs during sleep, suggesting that it may integrate environmental and cortical signals to regulate hippocampal network activity and gate sleep-dependent memory processing.

## Introduction

Animals continuously transition between behavioral states, including quiet wakefulness, heightened arousal, and sleep, in response to changing environmental and internal demands. This adaptability is essential for survival. State-dependent modulation of brain activity involves both ascending subcortical and descending cortical pathways and plays an important role in regulating neural processing, potentially enhancing or suppressing information flow among brain regions [1, 2]. Such flexibility enables the bidirectional modulation of neural circuit function and associated behaviors according to behavioral relevance.

Recently, key brain regions that critically regulate brain states have begun to be identified [2, 3]. One such region is the median raphe (MnR), located at the midbrain–pontine border [3–5]. Lesion or inactivation of the MnR induces robust behavioral activation, including increased locomotion, feeding, drinking, and gnawing [6, 7]. In contrast, stimulation of the MnR suppresses hippocampal–cortical activity and induces behavioral inhibition [4, 8, 9]. These findings indicate that the MnR plays a key role in regulating brain and behavioral states. Consistent with this notion, we have shown that many MnR neurons exhibit state-dependent activity across the sleep–wake cycle [4, 5]. However, how MnR neurons are regulated by distinct inputs across different brain states remains unclear. Both ascending subcortical and descending cortical pathways may contribute to the regulation of MnR neuronal activity, including during sleep, when the brain remains capable of processing diverse inputs to influence memory [10–12].

The MnR comprises three primary neuronal populations: serotonergic (5-HT), Vglut3-expressing, and GABAergic neurons [13–25]. More recently, Vglut2-expressing neurons have also been identified in the raphe region [26, 27]. However, this population appears to be predominantly restricted to the paramedian raphe and the caudal tail of the MnR [13, 27, 28], rather than the midline MnR. Among the three major MnR populations, 5-HT neurons exhibit clock-like activity that is largely independent of brain state [4, 5]. In contrast, Vglut3 neurons exhibit characteristic ON–OFF cyclic activity during sleep and may therefore contribute to state-dependent memory processes [4, 5], given their predominant projections to hippocampal network structures [19, 21, 24, 29]. GABAergic neurons, in turn, may regulate brain states through their projections to nearby Vglut3 neurons and also the interpeduncular nucleus [4, 5, 27, 29].

In the current study, we investigated how distinct MnR neuronal populations respond to sensory and prefrontal cortical inputs across wakefulness and sleep. We found that MnR neurons exhibited pronounced state-dependent responses to both sensory and cortical stimulation, with response amplitudes significantly greater during slow-wave sleep than during wakefulness.

## Results

### MnR neurons respond to sensory stimuli in a state-dependent manner

To identify genetically defined MnR Vglut3 neurons, we used the Vglut3-Cre transgenic mouse line [5]. Specifically, we employed an optogenetic tagging approach [5], involving intra-MnR microinjection of Cre-dependent AAV vectors encoding ChR2 (AAV-DIO-ChR2) and implantation of an optrode, consiting of an optical fiber surrounded by 4–6 tetrodes (Fig. 1A). After 2–3 weeks to allow AAV expression, we began recording the local field potential and spike activity of MnR neurons in freely behaving mice (Fig. 1B). Neurons that exhibited reliable, short-latency (3–10 ms) responses to blue-light pulse-train stimulation were classified as Vglut3 neurons (Fig. 1C). Consistent with previous findings [4, 5], these tagged Vglut3 neurons largely exhibited ON–OFF cyclic activity (∼50–100 sec per cycle) during slow-wave sleep (SWS; Fig. 1D).

**Figure 1.**
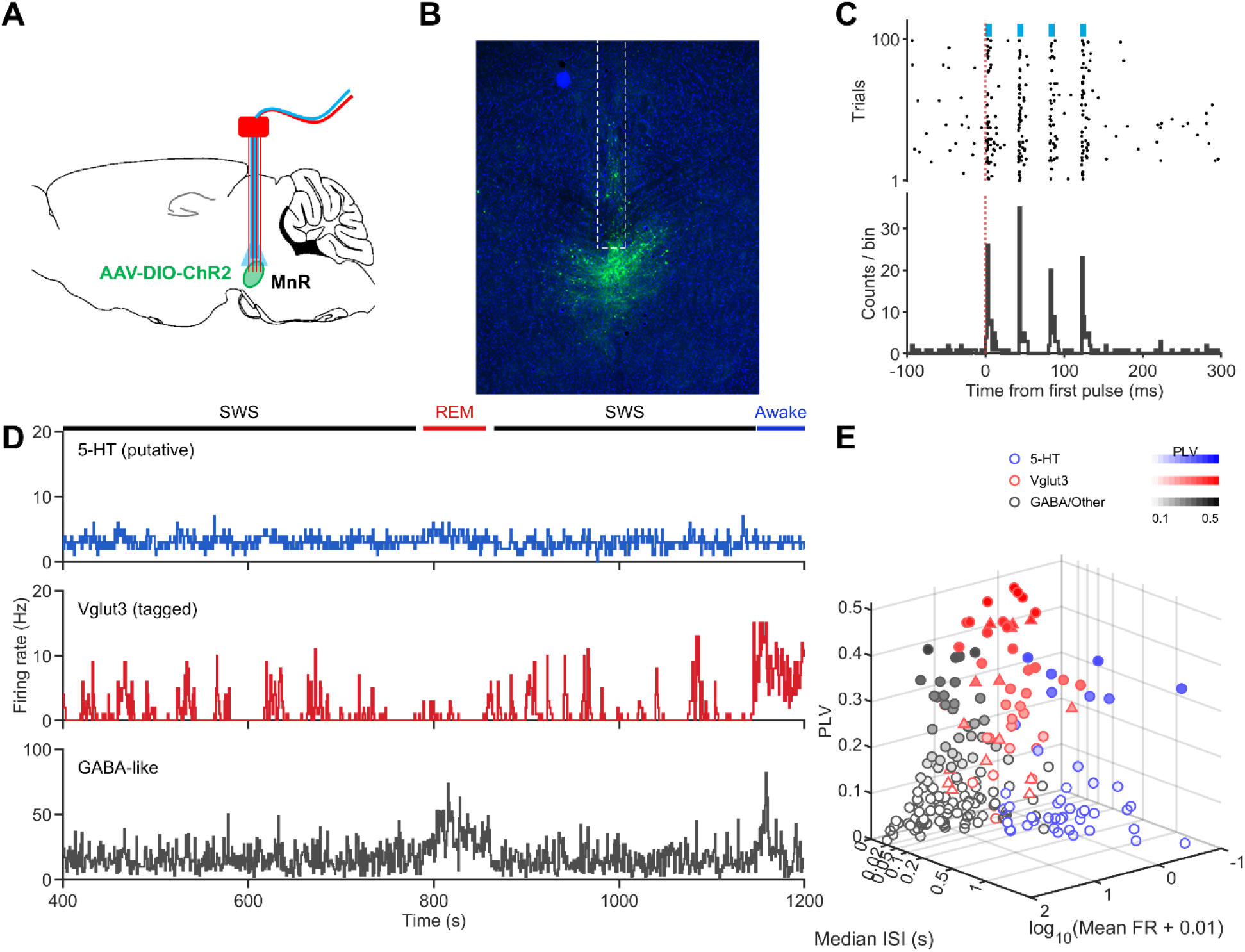
Three major MnR neuronal populations. **A,** Experimental design for optogenetic tagging. An optrode, consisting of an optical fiber surrounded by multiple tetrodes, was implanted into the MnR (injected with AAV-DIO-ChR2-EYFP) of Vglut3-Cre transgenic mice. **B,** Representative fluorescence image showing EYFP expression in MnR Vglut3 neurons. The white dashed rectangle indicates the optrode implantation track. **C,** Peristimulus rasters and histogram of a representative tagged Vglut3 neuron showing reliable firings upon optostimulation (4 pulses, 25 Hz; pulse width, 3 ms). **D.** Firing-rate histograms of three simultaneously recorded MnR neurons across sleep–wake states. The Vglut3 neuron is the same neuron shown in C, exhibiting a characteristic cyclic On–Off firing pattern during slow-wave sleep (SWS). REM, rapid eye movement sleep. **E.** Classification of MnR neurons based on mean firing rate (FR), median inter-spike interval (ISI), and phase-locking value (PLV; see Methods). 5-HT (blue), Vglut3 (red), and GABA/other (gray) neurons are shown, with triangles indicating optogenetically tagged Vglut3 neurons. Filled symbols indicate significant phase locking (p < 0.05), with color intensity reflecting the PLV.

The remaining MnR neurons were classified as either putative 5-HT or GABA/other neurons, the latter consisting primarily of GABAergic neurons and potentially some Vglut2 neurons. Putative 5-HT neurons were characterized primarily by clock-like firing patterns and long interspike intervals (median, ≥200 ms) [4, 5], whereas putative GABA/other neurons exhibited more diverse firing patterns (Fig. 1 D&E). Given the prominent ON–OFF cyclic activity pattern of Vglut3 neurons, we further classified a subset of putative Vglut3 neurons recorded in wildtype mice based primarily on the phase-locking of their activity to intrinsic infraslow fluctuations (0.01–0.02 Hz), together with additional firing characteristics (Fig. 1E; see Methods for details).

To determine how each type of MnR neuron responds to auditory stimuli across sleep–wake states, we continuously recorded MnR neuronal activity while intermittently delivering low-intensity white noise clicks (∼55 dB, 100 ms) with an inter-trial interval of 8–12 sec. The stimulus intensity was deliberately kept low to minimize auditory arousal and avoid disrupting the animals’ natural sleep–wake cycles. We found that all three MnR neuronal populations exhibited robust responses to the auditory stimuli during SWS, but their responses were substantially more muted during wakefulness (Fig. 2). Specifically, 5-HT neurons were largely suppressed, whereas both Vglut3 and GABA/other neurons were activated (Fig. 2). These findings suggest that MnR neurons exhibit robust and cell-type-specific responses to sensory stimuli during sleep.

**Figure 2.**
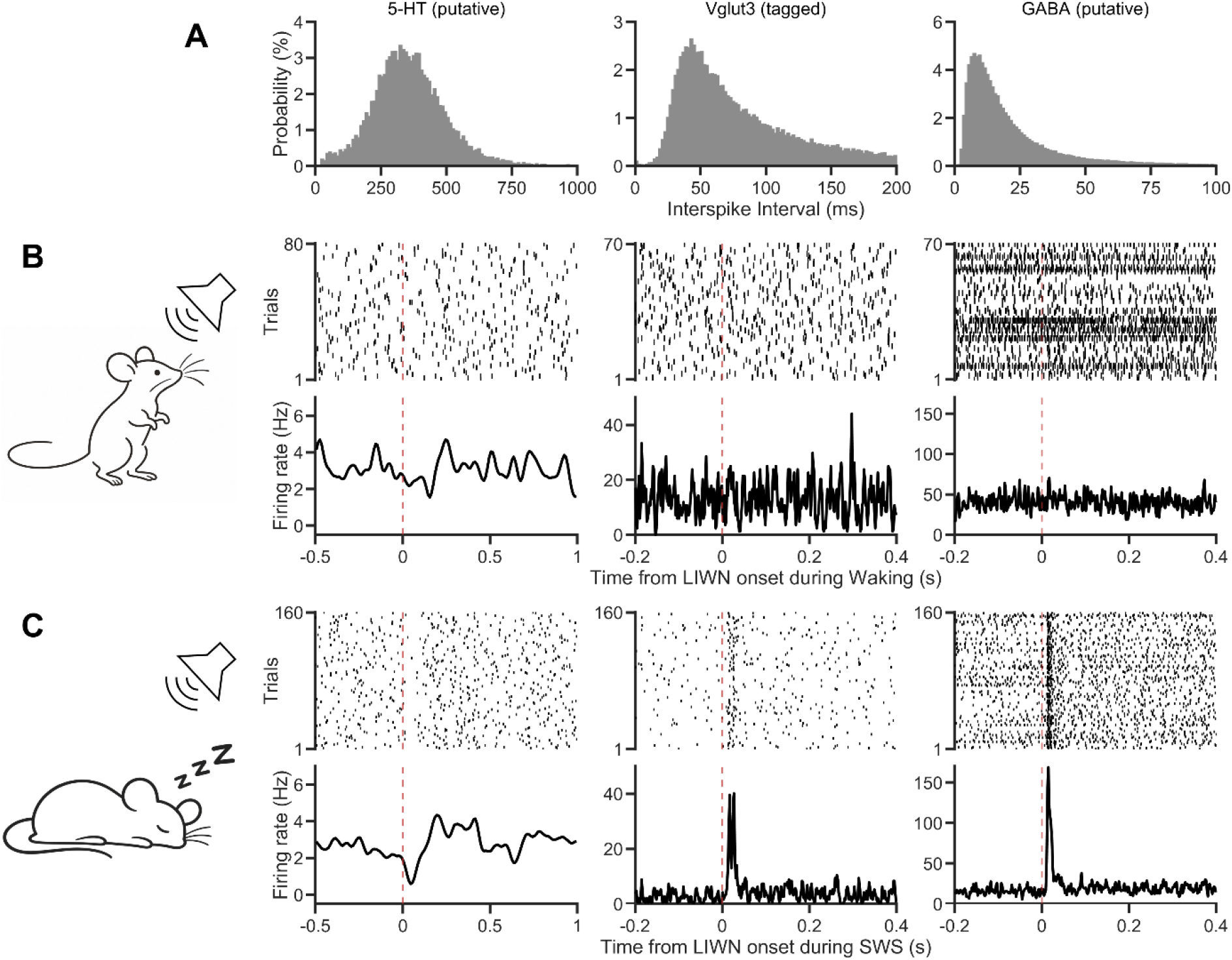
MnR neurons respond more strongly to non-arousal auditory stimuli during SWS than wakefulness. **A**, Representative inter-spike interval (ISI) distributions of three MnR neurons (putative 5-HT, tagged Vglut3, and GABA/other). **B**, Left, schematic of the experimental design for low-intensity white noise stimulation (LIWN, ∼55 dB, 100ms). Right, the three panels are peri-event rasters (top) and PSTHs (bottom) showing auditory responses of the same three MnR neurons during wakefulness. **C**, Schematic and responses of the same three MnR neurons to identical low-intensity auditory stimulation during SWS.

In total, we analyzed 42 putative 5-HT neurons, 27 Vglut3 neurons (12 tagged and 15 putative), and 137 GABA/other MnR neurons. During wakefulness, auditory stimulation elicited relatively modest responses across MnR neuronal populations (Fig. 3A). Among putative 5-HT neurons, 14.3% (6/42) were inhibited. In contrast, excitatory responses were observed in 40.7% (11/27) of Vglut3 neurons and 35.0% (48/137) of GABA/other neurons. During SWS, auditory-evoked responses became markedly more prevalent across all three neuronal populations (Fig. 3B). Most 5-HT neurons (52.4%, 22/42) exhibited suppression often followed by rebound activation. In contrast, Vglut3 and GABA/other neurons showed predominantly rapid excitatory responses, with 74.1% (20/27) and 79.6% (109/137) exhibiting excitation, respectively. Population-averaged firing activity further revealed more pronounced excitation in Vglut3 and GABA/other neurons during SWS than during wakefulness (Fig. 3C), highlightling their enhanced responsiveness to auditory stimulation during sleep.

**Figure 3.**
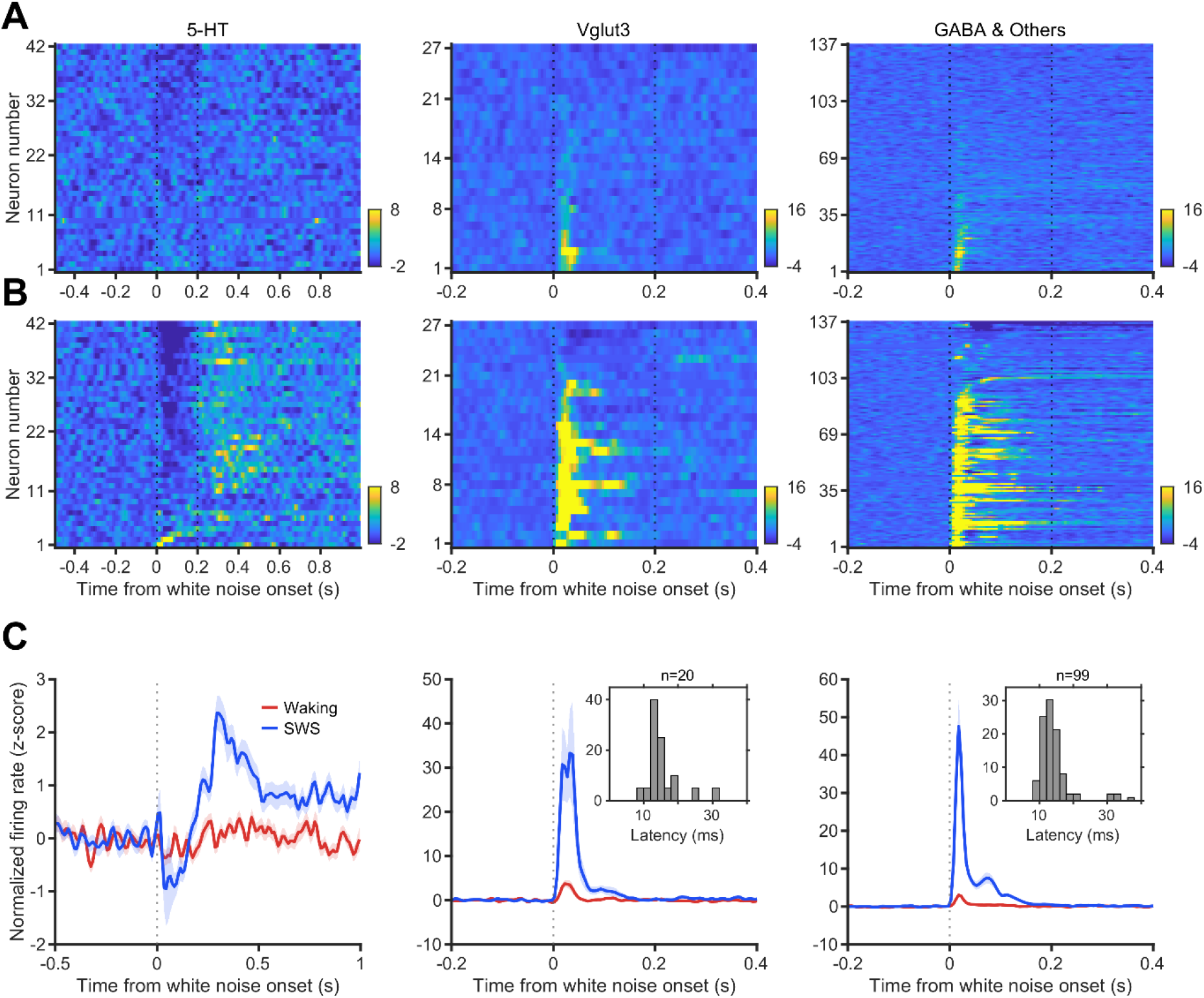
State-dependent responses of MnR neurons to non-arousal auditory stimulation. **A&B,** Heatmaps showing the normalized (z-scored) activity of 5-HT neurons (n = 42), Vglut3 neurons (n = 27), and GABA/other neurons (n = 137), aligned to the onset of a 100-ms low-intensity white-noise stimulus during wakefulness (A) and slow-wave sleep (SWS; B). Each row represents an individual neuron. Color indicates normalized activity relative to the pre-stimulus baseline. **C**. Mean (± SEM) normalized activity for each neuronal group during wakefulness (red) and SWS (blue). The 5-HT neurons exhibited auditory-evoked suppression followed by rebound activation during SWS, whereas Vglut3 and GABA/other neurons exhibited robust, rapid auditory-evoked excitation that was markedly greater during SWS than wakefulness. Insets show the distribution of response latencies for Vglut3 (middle) and GABA/other neuronal populations during SWS.

Together, these findings demonstrate a pronounced dependence of auditory responsiveness in the MnR on brain state. Compared with wakefulness, SWS was associated with more widespread auditory-evoked responses, characterized by predominantly inhibitory responses in 5-HT neurons and robust excitatory responses in Vglut3 and GABA/other neurons.

### MnR neurons exhibit short-latency auditory responses during SWS

To gain insight into the potential sources of auditory input to MnR neurons, we examined the onset latencies of sound-evoked responses. Because both Vglut3 and GABA/other MnR neurons exhibited prominent increases in firing, we quantified the onset latencies of their excitatory responses. Among neurons with detectable response within the first 40 ms after stimulus onset, Vglut3 neurons (20/27) exhibited a mean onset latency of 15.5 ± 5.0 ms (mean ± SD) and a median of 14 ms. Similarly, GABA/other neurons (99/137) exhibited a mean onset latency of 14.3 ± 5.0 ms and a median of 13 ms. In both populations, response onset latencies were concentrated between 10–20 ms (Fig.3C, inset panels). These findings indicate that auditory stimulation rapidly recruits both Vglut3 and GABA/other MnR neurons during SWS, suggesting potential contributions from relatively direct auditory pathways originating in the brainstem and/or thalamus.

### MnR neurons differentially respond to shake and visual stimulation

To determine whether MnR neuronal populations exhibit similar response patterns across sensory modalities, we further examined neuronal activity in response to brief earthquake-like shake [30] and light-flash stimulation, representing vestibular/somatosensory and visual inputs, respectively (Fig. 4). Shake stimulation elicited robust responses across MnR neuronal populations (Fig. 4A). Most 5-HT neurons (54.8%, 23/42) exhibited suppressed activity, whereas Vglut3 and GABA/other neurons showed predominantly excitatory responses, with 55.6% (15/27) and 62.8% (86/137) exhibiting excitation, respectively. These findings revealed distinct response patterns among MnR neuronal populations, consistent with those observed following auditory stimulation.

**Figure 4.**
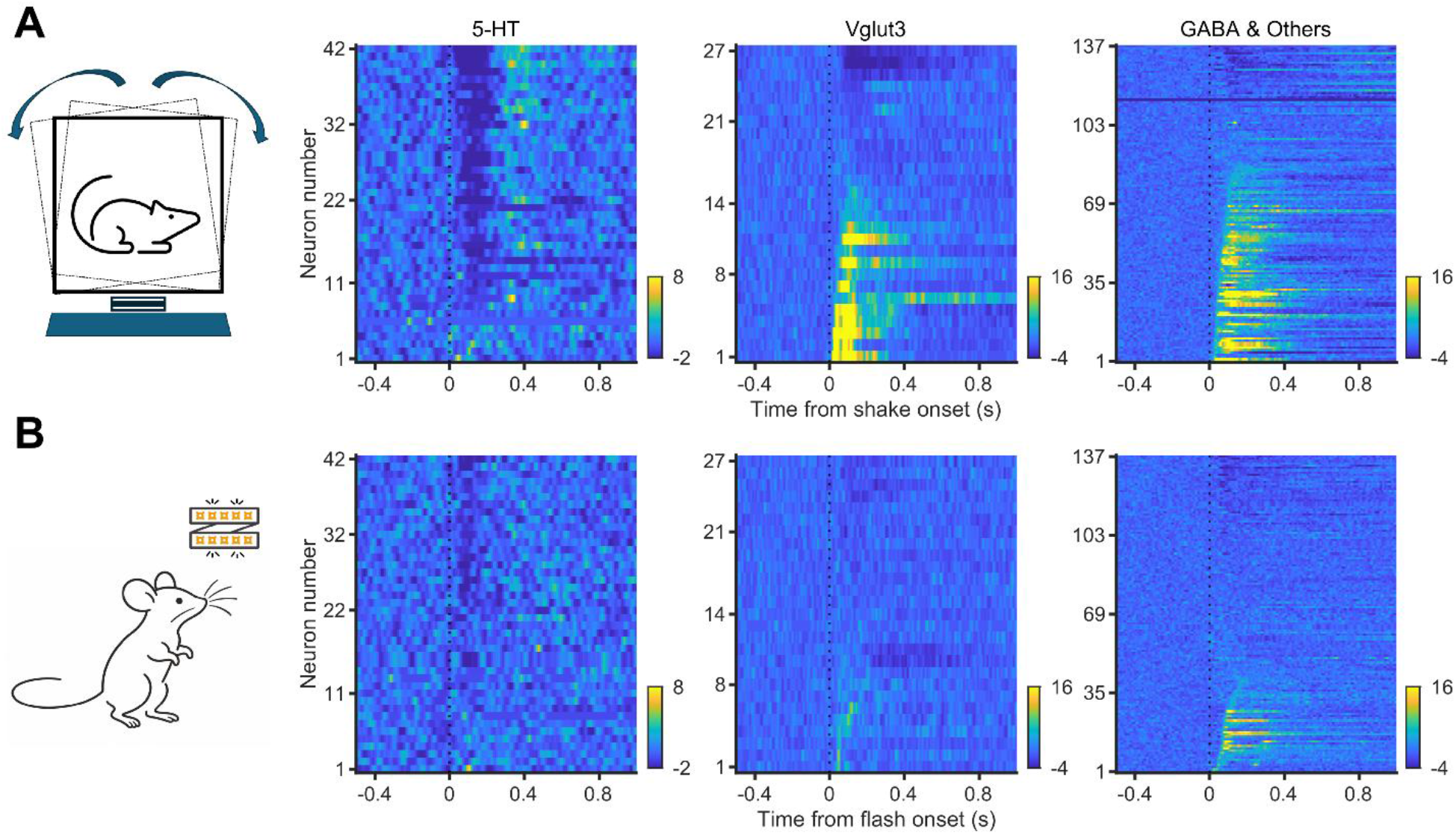
Responses of MnR neurons to shake and visual flash stimulation. **A,** Left, schematic of shake stimulation. Right, heatmaps showing the normalized (z-scored) activity of 5-HT (n = 42), Vglut3 (n = 27), and GABA/other neurons (n = 137), aligned to the onset of a brief shake stimulus. Each row represents an individual neuron. Color indicates normalized activity relative to the pre-stimulus baseline. **B,** Left, schematic of LED flash stimulation. Right, heatmaps showing the responses of the same neuronal populations to flash stimulation. Shake stimulation elicited prominent excitation in Vglut3 and GABA/other neuronal populations, whereas flash-evoked responses were weaker. The 5-HT population displayed prominent inhibitory responses to shake stimulation and weaker responses to flash stimulation.

Light-flash stimulation elicited a similar pattern of neuronal responses, although fewer Vglut3 and GABA/other neurons exhibited detectable responses (Fig. 4B). A subset of 5-HT neurons (33.4%, 14/42) was suppressed, whereas excitatory responses were observed in 29.6% (8/27) of Vglut3 neurons and 29.2% (40/137) of GABA/other neurons. Together, these findings suggest that MnR neuronal populations exhibit broadly consistent response patterns across sensory modalities, characterized by predominant suppression of 5-HT neurons and excitation of Vglut3 and GABA/other neurons.

### MnR neuronal response magnitudes are positively correlated across sensory stimuli

To further investigate whether MnR neurons exhibit shared response properties across sensory modalities, we examined pairwise correlations in neuronal responses to white noise, shake, and light flash stimulation during SWS (Fig. 5). For each neuron, excitatory and inhibitory responses were represented by positive and negative normalized peak z-scores, respectively. Neurons that did not exhibit detectable responses to either stimulus in a given pair were excluded from the corresponding comparison. Specifically, we quantified pairwise correlations in firing rate changes (ΔFR) within the 0–200 ms post-stimulus window for each sensory modality (Fig. 5).

**Figure 5.**
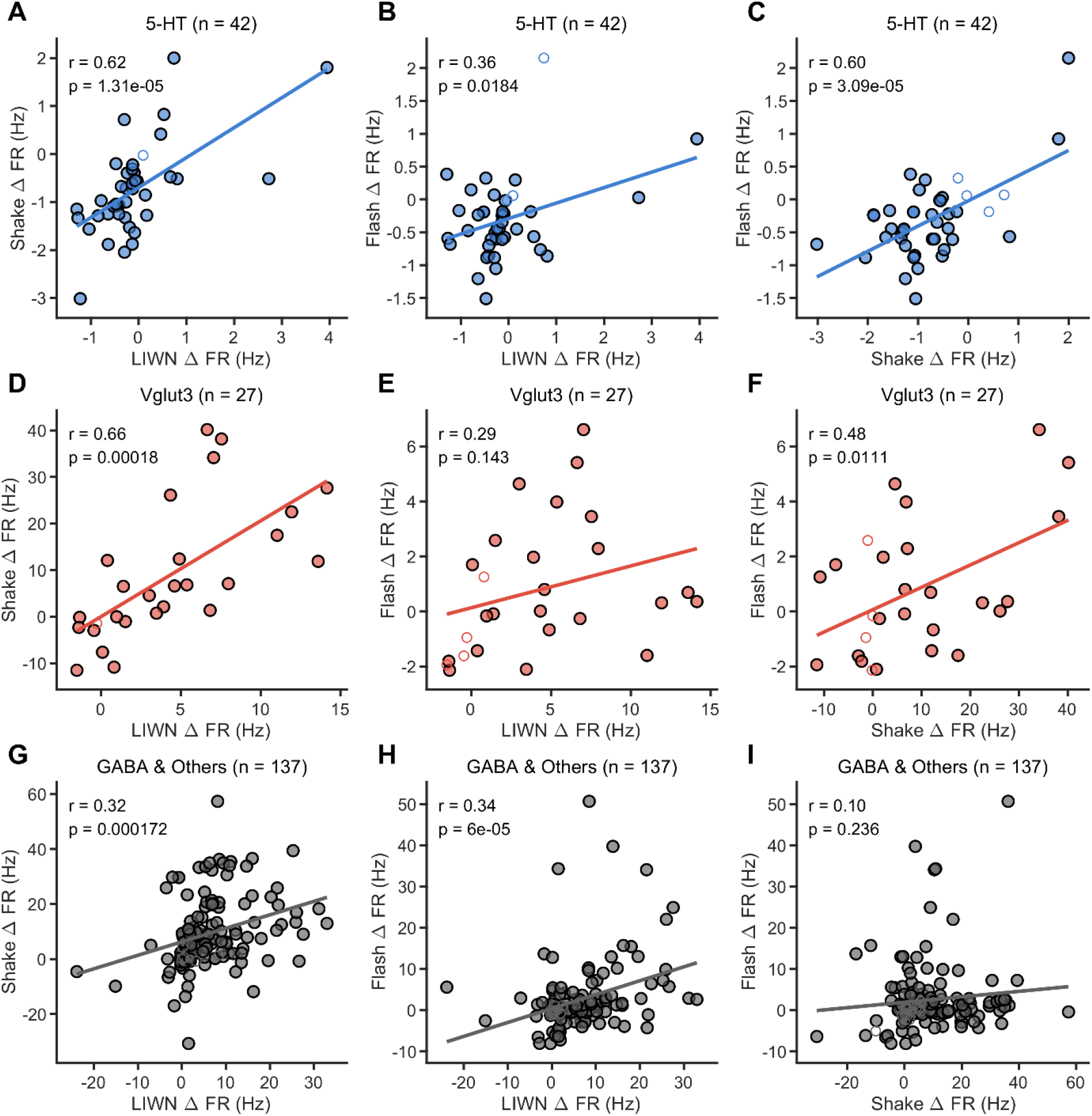
Correlations of sensory-evoked response magnitudes across MnR neuronal populations. **A–I,** Scatter plots showing pairwise correlations of firing rate changes (ΔFR) in response to low-intensity white noise (LIWN) during SWS, shake, and visual flash stimulation in individual MnR neurons. ΔFR was calculated as the difference between the mean firing rate during the 0–200 ms post-stimulus window and the baseline firing rate, expressed in Hz. Each dot represents an individual neuron. Neurons showing no detectable response to either stimulus in a given comparison are indicated by open circles, whereas the remaining neurons are shown by filled circles. **A–C,** Pairwise correlations in 5-HT neurons: LIWN versus shake (A), LIWN versus flash (B), and shake versus flash (C). **D–F,** Corresponding comparisons in Vglut3 neurons. **G–I,** Corresponding comparisons in GABA/other neurons. Solid lines indicate linear regression fits. Pearson correlation coefficients (r) and two-sided p values are indicated in each panel.

In 5-HT neurons, response magnitudes were positively correlated between white noise and shake stimulation (r = 0.618, p = 1.31 × 10⁻⁵), and between shake and flash stimulation (r = 0.596, p = 3.09 × 10⁻⁵). A weaker but significant correlation was also observed between white noise and flash responses (r = 0.362, p = 0.0184). Vglut3 neurons showed a similar pattern, with significant correlations between white noise and shake responses (r = 0.660, p = 0.00018) and between shake and flash response (r = 0.481, p = 0.0111), but not between white noise and flash (r = 0.289, p = 0.143). In GABA/other MnR neurons, white noise responses were correlated with both shake (r = 0.316, p = 0.00017) and flash responses (r = 0.336, p = 6.00 × 10⁻⁵), whereas shake and flash responses were not significantly correlated (r = 0.102, p = 0.236).

Together, these findings indicate that response magnitudes are generally positively correlated across sensory stimuli in MnR neuronal populations, although the strength and pattern of these correlations vary across neuronal subtypes. Notably, the consistent correlations between white noise and shake responses across all three populations suggest partially shared mechanisms underlying auditory- and shake-evoked esponses of MnR neurons.

### Retrograde tracing of inputs to the MnR

To determine major inputs to the MnR, we conducted intra-MnR microinjection of retrograde AAV vectors encoding fluorescent protein GFP (retroAAV-CAG-GFP [31]) (Fig. 6A). Four weeks after the injection, mice were perfused and their brains were processed for visulazation of retrogradely labeled neurons. We focused on forebrain, sensory thalamic, and brainstem structures, which are relatively less characterized. In the forebrain, dense labeling was observed primarily in the medial prefrontal cortex (mPFC; Fig. 6 B–D), including the anterior cingulate cortex, prelimbic and infralimbic cortex, and orbitofrontal cortex, in line with prior findings [32–35]. Within the brainstem, prominent labeling was found in the dorsal tegmentum of the midbrain and pons. Based on their stereotaxic location and cytoarchitectonic landmarks, labeled neurons were consistent with the laterodorsal tegmental nucleus (LDT) and adjacent dorsal pontine tegmental regions, including the nucleus prepositus hypoglossi (Fig. 6 E&F). Additional labeled neurons were observed in more caudal midline pontine and medullary reticular regions, notably the medial longitudinal fasciculus (MLF), suggesting that multiple brainstem nuclei send direct inputs to the MnR.

**Figure 6.**
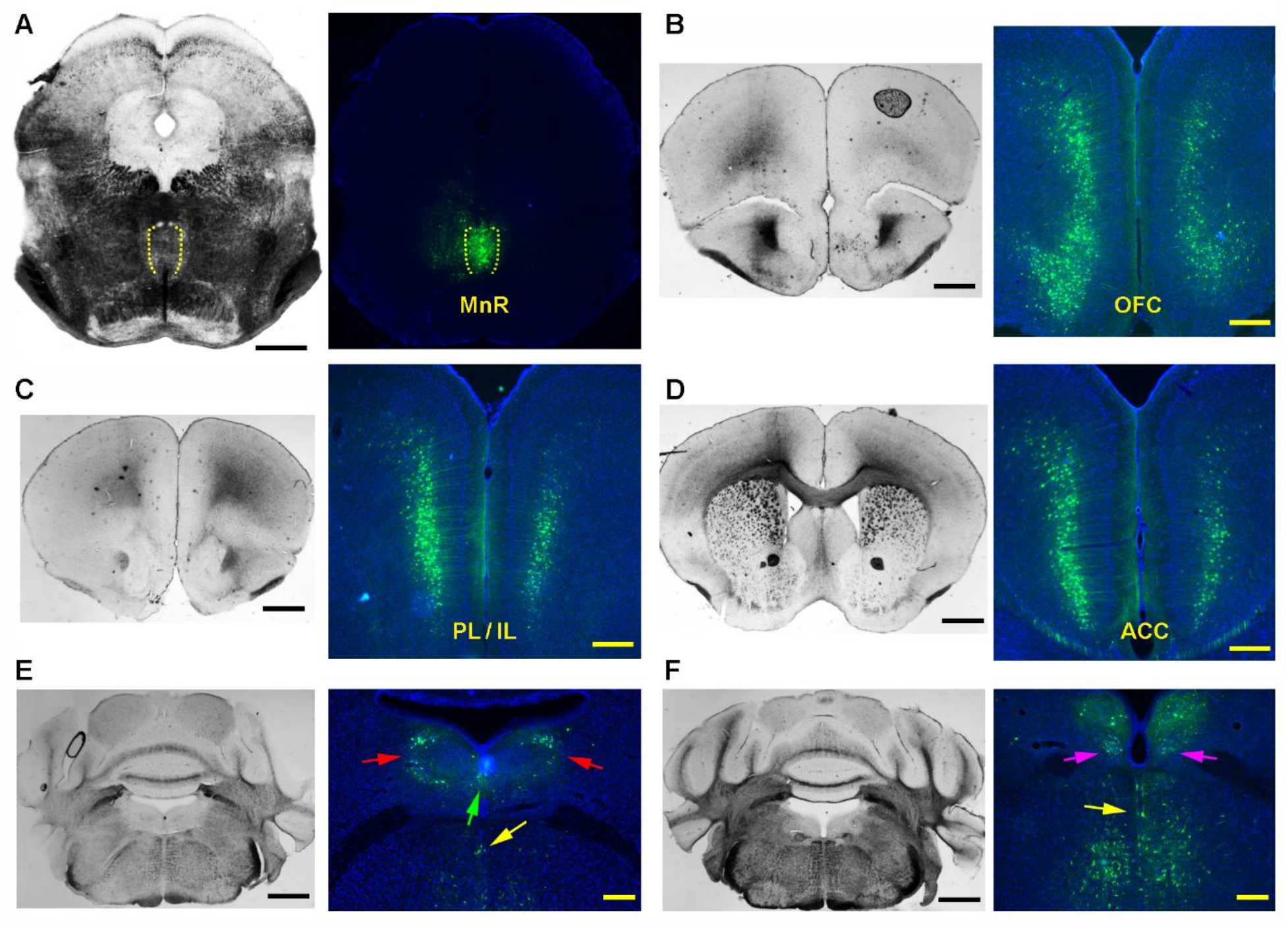
Retrograde tracing of cortical and brainstem inputs to the MnR. **A–F,** Bright-field images (left) and fluorescence images (right) centered on the MnR (A), orbitofrontal cortex (OFC; B), prelimbic (PL)/infralimbic cortex (IL; C), anterior cingulate cortex (ACC; D), tegmental nuclei (E&F), and the nucleus prepositus hypoglossi (F). This mouse received intra-MnR microinjection of retrograde AAV vectors encoding fluorescent protein GFP (retroAAV-CAG-GFP; 0.25 µL) and its brain was processed 4 weeks later. Note that the red, green, magenta, and yellow arrows indicate the laterodorsal tegmental nucleus (LDT), nucleus incertus (NI), nucleus prepositus hypoglossi (NPH), and medial longitudinal fasciculus (MLF), respectively. Scale bars, 1 mm.

### MnR neurons respond to mPFC inputs in a state-dependent manner

Given the prominent mPFC**→**MnR projection, we asked whether this pathway exhibits state-dependent regulation of MnR activity. To address this, we microinjected AAV vectors encoding ChR2 (AAV-CaMKII-ChR2) into the mPFC, centered on the anterior cingulate cortex. Approximately four weeks later, we implanted an optrode (an optic fiber surround by 6 tetrodes) into the MnR [4, 5]. We subsequently optogenetically stimulated mPFC neuronal axon terminals in the MnR while recording MnR spike activity cross the sleep–wake cycle (Fig. 7A). MnR neurons were classified as putative 5-HT, Vglut3, and GABA/other neuronal populations (see Methods). Optogenetic stimulation of mPFC terminals elicited distinct responses across these neuronal populations (Fig. 7B, C). Notably, most putative Vglut3 neurons exhibited excitatory responses (14/16 neurons, 87.5%), whereas a smaller proportion of GABA/other MnR neurons showed excitation (16/60 neurons, 26.7%). In contrast, most 5-HT neurons exhibited inhibitory responses (22/28 neurons, 78.6%). These findings suggest that mPFC inputs preferentially excite Vglut3 neurons in the MnR while exerting predominantly inhibitory effects on 5-HT neurons.

**Figure 7.**
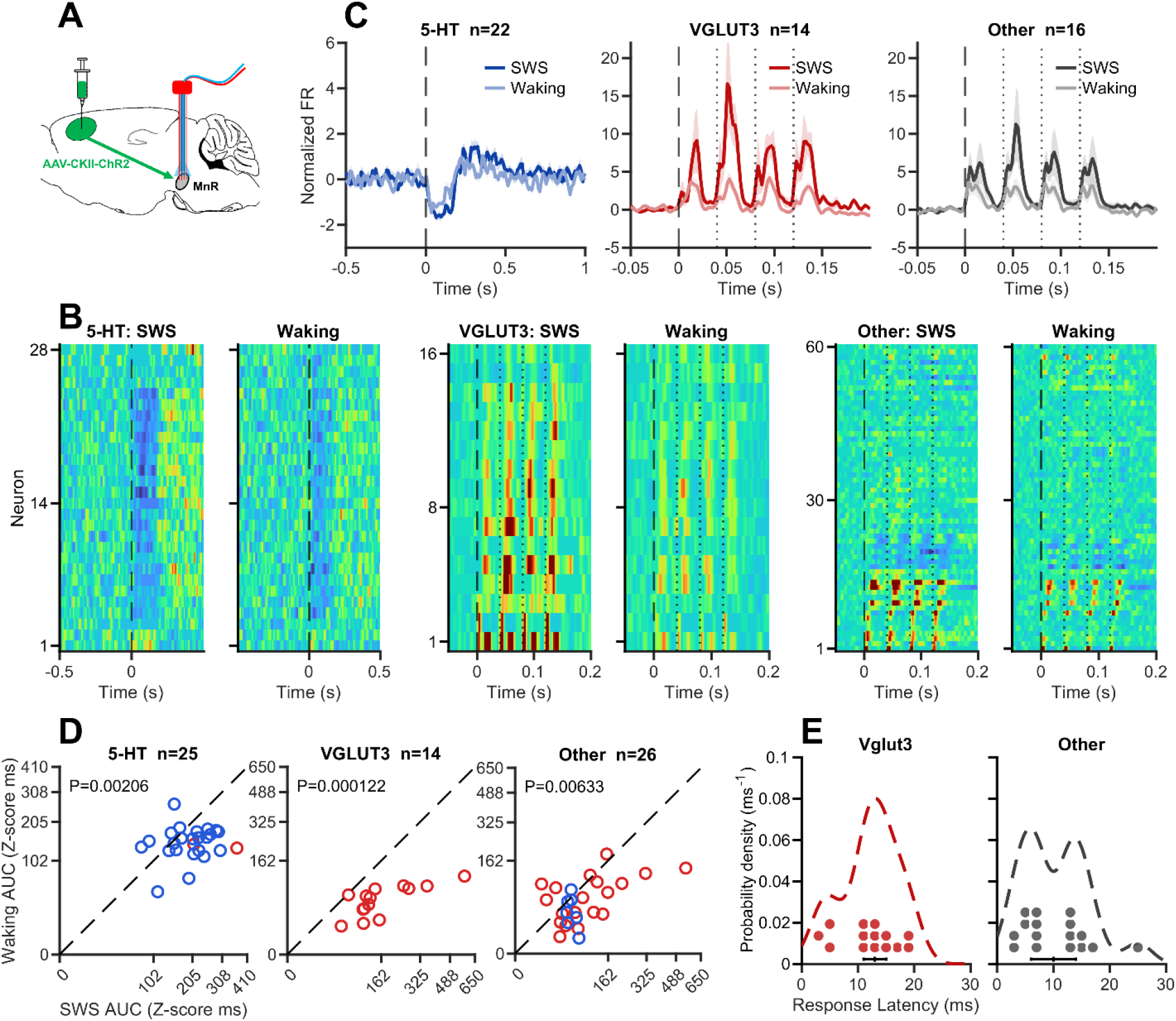
State-dependent responses of MnR neurons to mPFC inputs. **A,** Schematic of the experimental design: the mPFC received microinjection of AAV vectors encoding ChR2 (AAV-CaMKII-ChR2); the MnR received implantation of an optrode, consisting of an optical fiber surrounded by multiple tetrodes. This enabled recording of MnR neuronal activity upon optostimulation of mPFC neuronal axon terminals in the MnR (4 pulses at 25 Hz; pulse width, 10 ms). **B,** Heatmaps showing stimulus-aligned, baseline-normalized responses of putative 5-HT, Vglut3, and GABA/other neurons during SWS and wakefulness. Each row represents one neuron, with neurons arranged in the same order across states. **C,** Mean (± SEM) activity of the three MnR neuronal populations during SWS and wakefulness. Solid lines represent population means, and shaded areas indicate SEM. **D,** Paired comparisons of stimulation-evoked response magnitudes, quantified as the area under the curve (AUC) of normalized responses, between SWS and wakefulness. Each point represents one neuron. Response magnitudes were significantly greater during SWS in 5-HT (n = 25, Wilcoxon signed-rank test, P = 0.002), Vglut3 (n = 14, P = 0.0001), and GABA/other MnR neurons (n = 26, P = 0.006). **E.** Distributions of stimulus-evoked response latencies in excitatory-responsive Vglut3 (n = 14) and GABA/other neurons (n = 16) during SWS. Each dot represents one neuron, dashed curves indicate Gaussian kernel density estimates, and black horizontal bars indicate the interquartile range (IQR), with vertical ticks marking the median latency.

We next examined whether mPFC-evoked responses were modulated by brain state. Stimulation-evoked response magnitudes, quantified as the area under the curve (AUC) of normalized firing rate responses, were significantly greater during SWS than during waking across all three neuronal populations (Fig. 7D). This enhancement was observed in putative 5-HT neurons (P = 0.002; Wilcoxon signed-rank test), Vglut3 neurons (P = 0.0001), and GABA/other MnR neurons (P = 0.006). These findings indicate that mPFC-evoked responses of MnR neuronal activity is consistently enhanced during SWS across all three neuronal populations.

We further examined the temporal characteristics of mPFC-evoked excitation (Fig. 7E). Vglut3 neurons exhibited short-latency excitatory responses, with peak latencies ranging from 3–19 ms (median: 13 ms). Similarly, GABA/other MnR neurons exhibited short-latency responses, with peak latencies ranging from 3–25 ms (median: 10 ms). These rapid response latencies are consistent with effective synaptic drive of MnR neurons by mPFC inputs. Together, these findings suggest that mPFC terminal stimulation preferentially recruits Vglut3 neurons in the MnR, while the magnitude of stimulation-evoked responses is enhanced during SWS across all three neuronal populations. Thus, mPFC inputs exert both cell-type-dependent effects on MnR neuronal activity and state-dependent modulation of response magnitude.

## Discussion

### MnR neurons gate information flow in a state-dependent manner

Our results revealed that the three major populations of MnR neurons, including 5-HT, Vglut3, and GABAergic neurons, exhibited state-dependent responses to sensory and cortical inputs. Both sensory stimulation and optogenetic activation of prefrontal terminals elicited prominent responses in MnR neurons during slow-wave sleep, whereas responses were generally weaker during wakefulness. This state-dependent sensitivity of the MnR is notable because it contrasts with sensory processing in many brain regions, where neuronal responsiveness is typically enhanced during wakeful arousal [2, 36]. Nevertheless, sensory transmission is not completely suppressed during sleep. Sound-evoked responses remain detectable in the auditory midbrain, thalamus, and auditory cortex across vigilance states, although their magnitude and downstream propagation are strongly state-dependent [36–39]. Furthermore, sensory stimuli, including auditory and olfactory cues, can influence memory processes during sleep [10–12]. Together, these findings suggest that the MnR remains responsive to both sensory and cortical inputs during sleep and may serve as a state-dependent gate that regulates information processing in downstream brain regions.

MnR neurons, particularly Vglut3 neurons, project densely to the hippocampus and several other cortical regions involved in memory [19, 21, 24, 29, 40]. These MnR Vglut3 neurons may coordinate information flow in these interconnected regions through its predominant innervation of local GABAergic neurons. Within the hippocampus, MnR inputs selectively target non-parvalbumin GABAergic interneurons, while avoiding excitatory pyramidal neurons [24, 41–47]. Similarly, MnR inputs selectively synapse onto non-parvalbumin GABAergic interneurons in the frontal cortex [40, 48]. Therefore, state-dependent activation of MnR Vglut3 neurons may regulate hippocampal and cortical information processing through local inhibitory circuits, thereby gating information flow to memory-related regions.

The state-dependent sensory sensitivity of the MnR could be mediated by multiple mechanisms. The MnR may receive state-dependent sensory input. Several MnR-projecting brainstem nuclei, including the cochlear nucleus and superior colliculus, exhibit state-dependent sensory responsiveness similar to that observed in MnR neurons [49, 50]. It is also possible that MnR Vglut3 and GABAergic neurons are more excitable during SWS than during wakefulness. Increased membrane depolarization or reduced firing threshold during SWS would render these neurons more responsive to incoming excitatory inputs; however, future intracellular recordings will be required to directly test this possibility.

### MnR neurons receive sensory inputs from thalamic or brainstem regions

Previous work demonstrated that many MnR neurons respond to auditory stimuli with short latencies (6–12 ms) [51]. Similarly, neurons in nearby raphe nuclei can also respond to auditory or vestibular stimuli with short latencies (10–15 ms) [51–53]. Consistent with these findings, we observed that most sound-responsive MnR Vglut3 and GABAergic neurons responded between 10–20 ms during sleep, despite the very low intensity of the auditory stimuli used in our study (∼55 dB). These short latencies are comparable to the 12-ms auditory response latency reported for lateral amygdala neurons, which receive direct inputs from auditory thalamic regions [54]. Collectively, these findings suggest that auditory information can reach the MnR via relatively rapid pathways originating in lower-order thalamic or brainstem regions.

Indeed, the MnR is known to receive extensive inputs from pontine and medullary nuclei [27, 33, 55, 56], although many of these brainstem-afferents remain incompletely characterized. Our retrograde tracing analysis focused on brainstem regions, revealing dense labeling within dorsal pontine regions anatomically consistent with the laterodorsal tegmental nucleus (LDT) and neighboring pontine tegmental structures. Candidate auditory pathways may therefore involve the inferior colliculus together with the LDT, dorsal pontine tegmentum, or pontine reticular formation [27, 33, 55, 56]. However, the present anatomical data does not establish which of these nuclei conveys auditory information to the MnR or which MnR neuronal populations they directly target.

Overall, these findings indicate that short-latency auditory responses can arise through subcortical pathways without requiring extensive cortical processing. However, response latency alone cannot determine the exact number of synapses or identify the specific auditory pathways involved, as it is also influenced by stimulus intensity, axonal conduction velocity, and membrane integration [57]. Likewise, the heterogeneous excitation and inhibition observed among MnR 5-HT, Vglut3, and GABAergic neuronal populations may reflect distinct afferent pathways, recruitment of local neural circuits, or both.

### MnR neurons receive top-down prefrontal cortical inputs

In addition to sensory inputs, our results demonstrate that MnR neurons are functionally regulated by prefrontal cortical inputs. Optogenetic stimulation of mPFC axon terminals in the MnR elicited differential responses across MnR neuronal populations in a state-dependent manner, with responses more prominent during SWS than during wakefulness. The mPFC is involved in cognitive control, action monitoring, and behavioral adaptation to changing environmental demands [58–60]. Its projections to the MnR may therefore convey information about ongoing cortical activity and behavioral context. The enhanced responsiveness of MnR neurons to cortical stimulation during SWS suggests that mPFC inputs exert a stronger influence on MnR activity during this state, potentially contributing to sleep-related neural oscillations, information processing, and memory consolidation [61, 62].

The enhanced responsiveness to mPFC terminal stimulation during SWS also raises the possibility that cortical and sensory inputs converge on shared state-dependent mechanisms within the MnR. Such convergence may allow the MnR to integrate external sensory information with top-down cortical signals, thereby regulating the flow of information to downstream memory-related regions according to brain state. This mechanism may be particularly relevant during SWS, when MnR neurons exhibit heightened responsiveness to both sensory and cortical inputs, potentially facilitating the selective processing of information relevant to memory consolidation.

### Functional implications

Our findings suggest that the MnR participates in the state-dependent processing of cortical and sensory signals rather than acting as a simple relay of sensory information. The heterogeneous excitation and inhibition observed among distinct MnR neuronal populations further suggest that these inputs may be differentially processed within MnR circuits before influencing downstream targets. The functional significance of this state-dependent sensitivity may be twofold. During wakefulness, the relatively low responsiveness of MnR neurons to weak cortical and sensory inputs may limit unnecessary perturbations of ongoing neural activity. In contrast, during SWS, enhanced responsiveness may allow both cortical and external sensory signals to influence MnR activity and its downstream networks. In particular, the mPFC**→**MnR pathway may provide a mechanism through which cortical activity modulates subcortical circuits involved in hippocampal network regulation, whereas brainstem sensory pathways may convey information about environmental stimuli [10–12]. Although the present findings do not establish whether these inputs converge onto the same neurons or interact functionally, their shared state dependence suggests that the MnR may serve as an important site for coordinating cortical and sensory influences during sleep.

Recent work has demonstrated that non-arousing sounds presented during sleep influence hippocampal sharp-wave ripples and associated memory consolidation, indicating that sensory information continues to influence offline memory processing during sleep [12, 63]. Our results extend these findings by identifying the MnR as a potential upstream node through which sensory information may access the septohippocampal network. Because MnR Vglut3 neurons regulate hippocampal ripple activity and memory consolidation during sleep [4, 5], sensory activation of these neurons may provide a means of modulating hippocampal network activity without necessarily inducing behavioral arousal. Future studies combining closed-loop sensory stimulation with cell-type-specific manipulation of MnR neurons will be needed to determine whether this pathway causally links external sensory signals to hippocampal ripples and memory processes. Furthermore, integrating non-arousing sensory stimulation with top-down prefrontal inputs may provide a promising strategy for modulating memory processing during sleep.

## Materials and Methods

### Mice

Male C57BL/6J (Stock No. 000664) and Vglut3-Cre (Slc17a8-IRES2-Cre-D; Stock No. 028534) mice were purchased from the Jackson Laboratory and used for the experiments. Mice were 2.5–5 months old at the time of surgery. Following surgery, mice were individually housed in cages (40 × 20 × 25 cm) containing wood-chip bedding and a cardboard box filled with cotton nesting material. Mice were maintained on a 12 h light/dark cycle with ad libitum access to food and water. All experimental procedures were approved by the Institutional Animal Care and Use Committees of Drexel University (Protocol # LA-23-740) and adhered to the National Research Council’s Guide for the Care and Use of Laboratory Animals.

### Viral vectors

Adeno-associated viral (AAV) vectors, including AAV1-CaMKIIα-hChR2(H134R)-EYFP (#26973, Addgene), AAV1-EF1α-DIO-hChR2(H134R)-EYFP-WPRE-hGHpA (#20298), and retrograde retroAAV-CAG-GFP (#37825) were used for the experiments. Viral preparations were diluted to a final concentration of ∼5–10 × 10¹² genome copies/mL for microinjection.

### Stereotaxic surgery

Stereotaxic surgical procedures were performed as described previously [4, 5]. Briefly, mice were anesthetized with a ketamine/xylazine mixture (∼100/10 mg/kg, i.p.) and maintained on a heating pad at ∼37°C throughout surgery. For in vivo recording, both C57BL/6J and Vglut3-Cre mice received intra-MnR implantation of an optrode, consisting of an optical fiber surrounded by 4–6 tetrodes [4, 5]. In addition, Vglut3-Cre mice received intra-MnR microinjection of AAV-DIO-ChR2, whereas C57BL/6J mice received bilateral intra-mPFC microinjection of AAV-CaMKII-ChR2. To trace inputs to the MnR, C57BL/6J mice received intra-MnR microinjection of retrograde AAV (retroAAV-CAG-GFP).

The stereotaxic coordinates (Franklin and Paxinos 3^rd^ edition) relative to Bregma were as follows: MnR, anteroposterior (AP), −4.3 mm; mediolateral (ML), 0.0 mm; dorsoventral (DV), 3.9–4.0 mm; mPFC, AP, +1.2 mm; ML, 0.3 mm; DV, 1.2 mm. AAV vectors (0.2–0.25 μL) were delivered using a syringe pump (World Precision Instruments) at a rate of 0.05 μL/min. Following each injection, the injection needle (34 gauge, beveled) was left in place for an additional 5–10 min to minimize viral reflux before being slowly withdrawn. Optrodes were gradually lowered into the MnR and secured to the skull using stainless-steel screws and dental acrylic. Electrophysiological recordings and optogenetic experiments began 2–3 weeks after surgery to allow sufficient time for AAV expression.

### Tetrodes and optrodes

Each tetrode consisted of four platinum–iridium wires (90% platinum/10% iridium; 18 μm in diameter; California Fine Wire), with an impedance of ∼1–2 MΩ per wire. Each optrode consisted of an optical fiber (105 μm core diameter) surrounded by 4–6 tetrodes [4, 5]. The optical fiber tip was positioned ∼0.3–0.6 mm from the tetrode tips. Optrodes were mounted on custom microdrives, allowing gradual adjustment of recording depth across sessions [4,5]. Optogenetic stimulation was delivered at a light power of ∼0.5–2 mW, with variable inter-stimulation intervals of 8–12 s.

### In vivo electrophysiology

Extracellular neural signals were amplified, digitized, and recorded using CerePlex acquisition system (Blackrock Neurotech). Animal behavior was simultaneously monitored and video-recorded during electrophysiological recordings. Local field potential (LFP) signals were sampled at 2 kHz and low-pass filtered at 500 Hz, with ground serving as the reference. Extracellular spike signals were sampled at 30 kHz and band-pass filtered at 600–6000 Hz. A recording electrode without detectable spike activity was selected as the reference for spike recordings. Following each recording session, the optrode was advanced by ∼50–100 μm to sample additional neuronal populations. Each mouse underwent ∼3–6 recording sessions, with individual sessions lasting up to 6 hours.

### Histological verification of optrode placement

At the completion of electrophysiological recordings, the final electrode positions were marked by passing a 10-μA current for 20 s through tetrodes using a linear constant-current stimulus isolator (NeuroLog System). Mice were then deeply anesthetized and transcardially perfused with ice-cold PBS or saline, followed by 10% formalin. Brains were removed and post-fixed in formalin for at least 24 h. Following fixation, brains were sectioned coronally at a thickness of 50 μm using a vibratome (Leica). Sections were mounted with Mowiol mounting medium mixed with DAPI (Vector Laboratories) for fluorescence microscopy to verify AAV expression and optrode placements.

### Electrophysiological data analysis

To ensure that the recorded neurons mainly originated from the MnR, only sessions containing 5-HT or Vglut3 neurons (either optogenetically tagged or putatively identified) were included in the analysis. A total of 104 MnR neurons were recorded from C57BL/6J mice (n = 5), and 102 MnR neurons were recorded from Vglut3-Cre mice (n = 8). Individual neuronal units were isolated based on extracellular spike waveforms using Offline Sorter (Plexon Inc.). Spike sorting incorporated multiple waveform features, including principal component analysis and spike energy, to distinguish individual neuronal units. Sorted spike trains were subsequently processed and analyzed using NeuroExplorer 5 (Nex Technologies) and custom-written scripts in MATLAB (MathWorks). Neurons with mean firing rates <0.2 Hz were excluded from subsequent analyses.

### Sensory stimulation

#### Low-intensity auditory stimulation

To characterize the responses of MnR neuronal populations to auditory stimuli across sleep–wake states, spike activity was continuously recorded while brief, low-intensity white-noise (LIWN) stimuli (∼55 dB, 100 ms duration) were delivered intermittently at variable inter-stimulus intervals of 10–15 s. The relatively low stimulus intensity was selected to minimize disruption of spontaneous sleep–wake states while allowing assessment of state-dependent neuronal responses. Auditory stimuli were delivered through a speaker positioned ∼50 cm above the home cage.

#### Other sensory stimulation

To investigate whether MnR neuronal populations respond to sensory inputs across different modalities, we additionally examined neuronal responses to brief visual (light-flash) and vestibular/somatosensory (earthquake-like shake) stimuli. Visual stimuli were delivered using LED lights mounted on the ceiling ∼50 cm above the home cage. Shake stimulation was generated using a custom-built shaking apparatus consisting of a black cylindrical chamber mechanically coupled to a vortex mixer [30]. Brief activation of the vortex mixer produced rapid orbital vibrations, generating a reproducible earthquake-like shaking stimulus. All sensory stimuli were controlled by a Master-8 pulse generator (A.M.P.I.) and triggered by 100-ms electrical pulses.

### Identification of SWS and REM sleep

Sleep–wake states were identified based on local field potential (LFP) activity recorded from the MnR in conjunction with simultaneous video monitoring of animal behavior, as previously described [4, 5]. SWS and REM sleep were distinguished based on the ratio of theta (6–10 Hz) to delta (1–4 Hz) power derived from LFP spectrograms. A theta/delta power ratio ≥2 was classified as REM sleep, whereas a ratio ≤1 was classified as SWS [4]. These electrophysiological criteria were combined with video recordings to confirm wakefulness, SWS, and REM sleep.

Sleep episodes were typically observed when mice remained immobile in a cardboard box containing cotton nesting material within their home cage, which served as their preferred sleeping area [4, 5]. Each REM episode was preceded by an SWS episode, and mice typically exhibited brief movements or awakened at the end of REM sleep. All C57BL/6J and Vglut3-Cre mice exhibited clearly distinguishable SWS–REM sleep cycles during recording sessions.

### Classification of MnR neurons

MnR neurons were classified based on their electrophysiological properties, including mean firing rate, inter-spike interval (ISI) distribution, and sleep-state-dependent firing patterns, together with optogenetic tagging when available. Mean firing rates were calculated by dividing the total number of spikes recorded during wakefulness and sleep by the combined duration of both states. ISIs were calculated from consecutive spike timestamps across the entire recording, and the median ISI was used to characterize the typical firing interval of each neuron.

Putative 5-HT neurons were identified based on their low firing rates (mean, ≤4 Hz) and regular, clock-like firing patterns, characterized by relatively long ISIs (median, ≥200 ms) [4, 5]. In addition to optogenetically tagged Vglut3 neurons (Fig. 1C), putative Vglut3 neurons were identified based primarily on their characteristic ON–OFF cyclic firing pattern during SWS, consisting of alternating periods of neuronal activation and suppression (Fig. 1D). Specifically, the phase-locking value (PLV) of each MnR neuron relative to the ON–OFF cycle was calculated (see below). Neurons with a PLV ≥0.1, a mean firing rate ≤12 Hz, and a median ISI ≥25 ms were classified as putative Vglut3 neurons (Fig. 1E). Finally, neurons that could not be assigned to either the 5-HT or Vglut3 population were categorized as GABA/other neurons.

To quantify phase locking to intrinsic infraslow firing-rate fluctuations (0.01–0.02 Hz) during SWS, continuous SWS episodes lasting ≥200 s were analyzed separately. Within each episode, spike timestamps were randomly divided into two approximately equal subsets (A and B). The firing rate of subset A was calculated using 1-s bins, smoothed with a 5-s Gaussian window, and bandpass-filtered at 0.01–0.02 Hz using a third-order Butterworth filter with zero-phase filtering. The instantaneous phase, *φ_A_*(*t*), was extracted using the Hilbert transform:

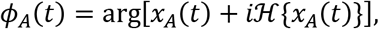

where *x_A_*(*t*) represents the filtered firing-rate signal and ℋ denotes the Hilbert transform. The instantaneous phase at each spike time in subset B was obtained by interpolating the complex phase representation, *e^iφA^*^(^*^t^*^)^. The procedure was then repeated with the subsets exchanged, using subset B to estimate the reference phase and subset A for spike-phase sampling. Complex phase vectors from both directions were pooled across valid SWS episodes, and the phase-locking value (PLV) was calculated as

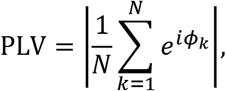

where *φ_k_* is the reference phase assigned to the *k*th held-out spike and *N* is the total number of valid spike-phase samples. PLV ranges from 0 to 1, with higher values indicating stronger phase concentration. To minimize filtering edge artifacts, 40 s were excluded from each end of each SWS episode. Statistical significance was assessed using 500 circular-shift surrogates, in which held-out spike times were randomly shifted relative to the reference phase series by at least 30 s within each valid analysis interval. The empirical *p* value was calculated as

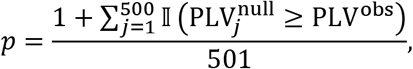

where I denotes the indicator function. Phase locking was considered significant at *p* < 0.05. For non-5-HT neurons, at least 1,000 spikes across all annotated SWS intervals were required for phase-locking significance analysis; neurons below this threshold were classified as nonsignificant. In Fig. 1E, neurons exhibiting significant phase locking were displayed with filled markers, with color intensity scaled according to PLV (0.1–0.5), whereas nonsignificant neurons were represented by open markers.

### Analysis of neuronal responses to sensory and optogenetic stimulation

Neuronal responses to sensory and optogenetic stimulation were analyzed using peri-event time histograms (PETHs) aligned to stimulus onset. For sensory stimulation, spike trains were binned at 5 ms for Vglut3 and GABA/other neurons and 10 ms for 5-HT neurons, with five-bin Gaussian smoothing and baseline periods of −0.5 to 0 s and −1 to 0 s, respectively. For mPFC**→**MnR terminal stimulation, corresponding bin sizes were 2 and 10 ms, smoothing windows were five and three bins, and baseline periods were −1 to 0 s and −5 to 0 s, respectively. Firing rates were normalized to baseline activity using Z-scores.

For sensory stimulation, neuronal responses were classified within 0–200 ms after stimulus onset using Z-score thresholds of +3/−2, requiring three consecutive bins exceeding the corresponding threshold. Stimulus-evoked changes in firing rate (ΔFR) were calculated as the difference between the mean firing rate during the 0–200 ms post-stimulus window and baseline. Response onset latency was defined as the first of three consecutive bins exceeding the corresponding Z-score threshold. For LIWN stimulation, onset latencies were calculated from independently generated PETHs using 2-ms bins for Vglut3 and GABA/other neurons and 10-ms bins for 5-HT neurons, with three-bin Gaussian smoothing. Onset latency distributions were analyzed within 0–40 ms after LIWN onset for Vglut3 and GABA/other neurons classified as excitatory responders during SWS.

For mPFC**→**MnR terminal stimulation, neuronal responses were classified as excitation, inhibition, or no response based on SWS activity. Responses were examined within 0–40 ms after individual light pulses for Vglut3 and GABA/other neurons and within 0–150 ms after the first pulse of each train for 5-HT neurons. Excitation and inhibition were identified using Z-score thresholds of +3/−2 for Vglut3 and GABA/other neurons, requiring a qualifying response peak or trough spanning three bins, and mean Z-score thresholds of +1/−0.5 for 5-HT neurons. Peak latency was defined as the time of the first qualifying response peak. Peak latency distributions were examined for SWS-excited Vglut3 and GABA/other neurons. Response magnitude was quantified as the direction-specific area under the Z-score curve (AUC) to compare SWS and wakefulness, including neurons responsive in either state.

### Statistical analysis

Sample sizes were determined based on previous studies using similar experimental approaches in our laboratories [4,5]. Statistical analyses were performed using MATLAB R2023b and SPSS Statistics 20.0 (IBM). Differences in optogenetically evoked response magnitudes between SWS and wakefulness were assessed using two-sided paired Wilcoxon signed-rank tests. Relationships between neuronal responses to different sensory modalities were assessed using Pearson correlation analyses. Spearman rank correlations were also calculated, and linear regression was used for visualization. All neurons with valid paired measurements were included regardless of response classification. Data are presented as mean ± SEM unless otherwise specified. Statistical significance was defined as p < 0.05, with additional statistical procedures indicated in the corresponding figure legends.

## Author contributions

W.H. and D.V.W. conceived and designed the experiments and wrote the manuscript. W.H. performed the experiments and analyzed the data.

## Financial Disclosures

The authors declare no financial interests or potential conflicts of interest.

## Acknowledgements

This work was supported by the National Institutes of Health grant R01MH119102.

## Notes

### Competing Interest Statement

The authors have declared no competing interest.

